# Self-sensitisation of *Staphylococcus aureus* to the antimicrobial factors found in human blood

**DOI:** 10.1101/2021.07.19.452914

**Authors:** Edward Douglas, Tarcisio Brignoli, Mario Recker, Eoin O’Brien, Rachel M. McLoughlin, Ruth C. Massey

## Abstract

For opportunistic pathogens, the switch from a commensal to an invasive lifestyle is often considered an accidental event. But with plentiful opportunity, what leads one accidental event to result in an invasive infection, and another not to? And how much of this apparent stochasticity is driven by bacterial factors? To answer these questions, here we focussed on the major human pathogen *Staphylococcus aureus,* which can both reside asymptomatically as a member of our respiratory microbiome, or become invasive and cause infections as severe as bacteraemia. Survival upon exposure to the antibacterial factors found in serum is a critical aspect of their ability to cause bacteraemia, and across a collection of 300 clinical isolates we found there to be significant variability in this capability. Utilising a GWAS approach we have uncovered the genetic basis of much of this variability through the identification and functional verification of a number of new polymorphic loci that affect serum survival: *tcaA, tarK, gntR, ilvC, arsB, yfhO,* and *pdhD.* The expression of one of these genes, *tcaA,* was found to be induced upon exposure to serum, while simultaneously enhancing the sensitivity of *S. aureus* to serum through a process involving the ligation of wall teichoic acids into the cell wall. As blood-stage infections are a transmission dead-end for the bacteria, that *S. aureus* actively responds to serum to produce a protein which specifically limits their ability to survive in this environment demonstrates that the switch from the commensal to the invasive lifestyle is complex, and that TcaA may contribute to the long-term success of *S. aureus* by restricting the bacteria to their more readily transmissible commensal state.

## Main Text

*Staphylococcus aureus* is an important human pathogen and a significant global health concern^1,2^. The most common interaction with its human host is as an asymptomatically coloniser; however, it frequently transitions to a pathogenic state with the ability to cause a wide range of diseases, ranging from relatively minor skin and soft tissue infections (SSTI) to more life-threatening incidents of endocarditis or bacteraemia^1,2^. *S. aureus* is notorious for producing a plethora of virulence factors ranging from pore forming toxins to various immune evasion strategies^3^. The toxicity of *S. aureus* is generally accepted as playing an important role during infection, with high toxicity isolates typically causing more severe symptoms and disease progression^4,5^. However, for the more invasive diseases such as bacteraemia and pneumonia, it has been shown that the causative isolates are often impaired in their toxin production^6,7^ and instead rely on alternative virulence approaches to cause disease, such as being able to better survive exposure to host immune defences.

As the most severe type of infection caused by *S. aureus,* bacteraemia has a mortality rate of between 20 and 30%^8^. Entry into the bloodstream is the first step in the development of bacteraemia and represents a major bottle neck for the bacteria, whether they seed from an infection elsewhere in the body, or they take a more direct route through an intravenous device. As a heavily protected niche, the bloodstream contains numerous humoral immune features with potent anti-staphylococcal activity, including antimicrobial peptides (AMPs) and host defence fatty acids (HDFAs)^9–12^. However, the fact that cases of *S. aureus* bacteraemia occur demonstrates that these do not represent an impregnable force and that the bacteria can adapt to resist these defensive features. For cationic AMPs, resistance is typically achieved through changing the charge of the outer cell envelope by the *dlt* operon^13^, or the membrane through the action of *mprF*^14^, such that the positively charged AMPs are partially repelled. *S. aureus* also contains numerous efflux pumps, sensing systems and can even secrete extracellular proteases to cleave the AMPs^15^. For HDFAs, resistance is less well characterised, but appears to be achieved by increasing the hydrophilicity of the cell envelope which impairs the penetration of the highly hydrophobic HDFAs^16^. While our understanding of the development of *S. aureus* bacteraemia is growing, two fundamental biological question remain: (1) as transmission rarely occurs from cases of bacteraemia, rendering it an evolutionary dead-end, why are the bacteria so well equipped for life in the bloodstream? And (2) given how well equipped the bacteria are for life in the bloodstream, why isn’t the incidence of *S. aureus* bacteraemia higher?

In previous work we have pioneered the application of genome wide association studies (GWAS) to characterise bacterial virulence, where it has proven to be a powerful approach to define complex regulatory pathways^7, 17–21^. In related work we have also begun to characterise a new category of *S. aureus* genes we refer to as MALs (mortality associated loci) and found that many of these are involved in serum survival^22^. Given the outstanding research questions surrounding the development of bacteraemia, and critical importance of serum resistance to patient outcome, we performed a GWAS on a collection of 300 clinical isolates with respect to serum resistance. We found significant variability in how well individual isolates survive exposure to serum and identified seven novel effectors of this activity. Of particular note was the TcaA protein, the expression of which we found to be both activated in response to serum while simultaneously sensitising the bacteria to serum. With such an unexpected activity, we propose that TcaA may provide a means by which *S. aureus* limits its ability cause bacteraemia and as such may play a key role in the long-term success of *S. aureus* by preventing the bacteria from entering into the evolutionary dead-end presented by the bloodstream.

## Results

The human bloodstream is a highly protected niche, and so to establish an infection in this environment the bacteria must evade many aspects of host immunity, such as the bacterial membrane damaging antimicrobial peptides (AMP) and host defence fatty acids (HDFAs) found in serum^9–15^. To examine the level of variability that exists in serum-susceptibility in natural populations of *S. aureus,* we exposed 300 clinical isolates to human serum and quantified the proportion of each culture that survived. These 300 isolates represent two major clones as defined by multi-locus sequence typing: clonal complex 22 (CC22) and clonal complex 30 (CC30)^18,21^. There were similar levels of variability across the two CCs, both with a >10^3^ fold difference between the most and least susceptible isolate with respect to the number of bacteria that survived exposure to serum (Fig. 1a&b).

**Figure 1:**
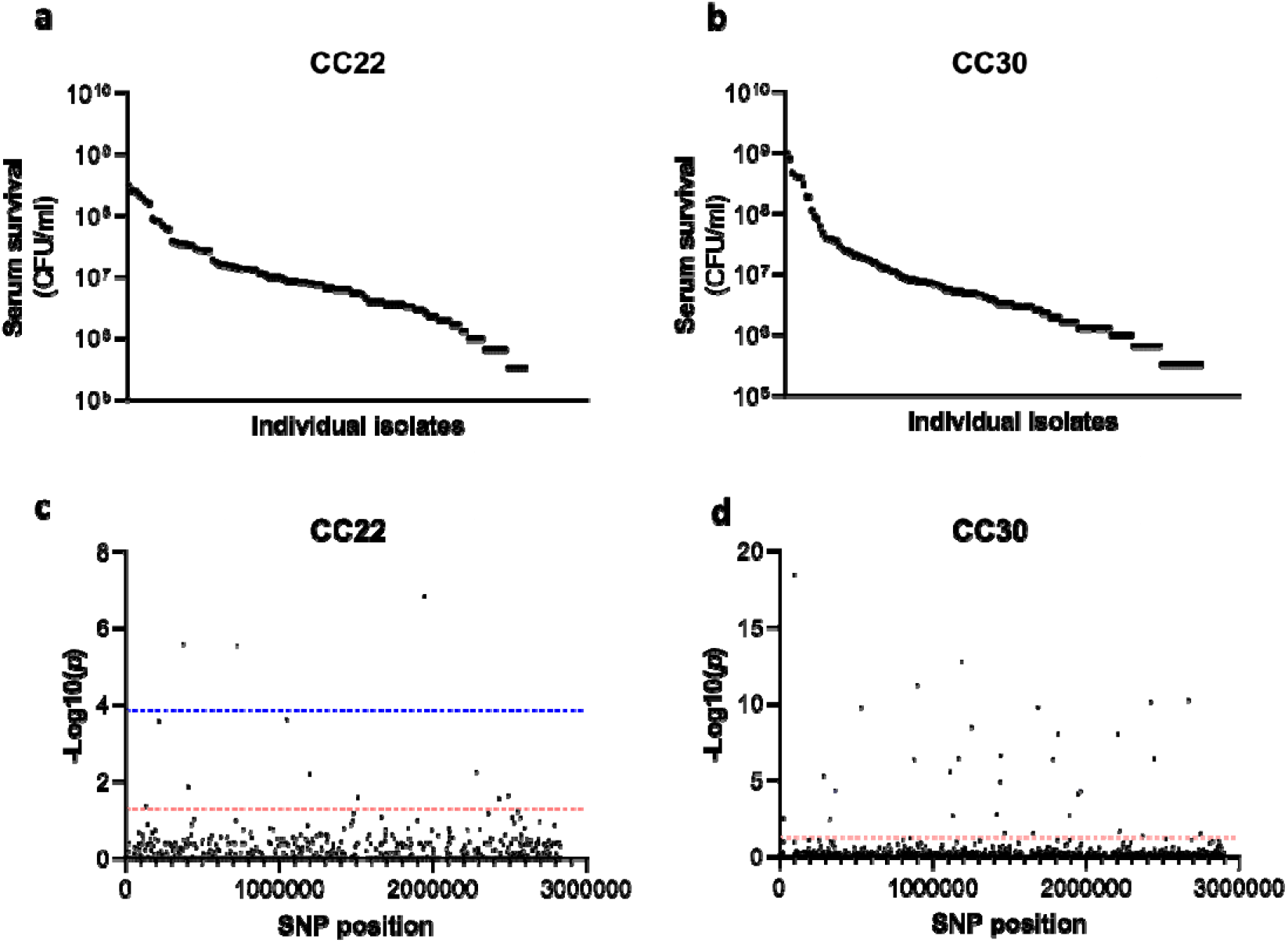
Serum susceptibility is multifactorial and varies widely across closely related *S. aureus* isolates. (**a & b**) The susceptibility of 300 bacteraemia isolates from clonal complexes CC22 and CC30 were exposed to human serum and their ability to survive this exposure quantified. Each dot represents the mean survival of an individual isolate. (**c & d**) Manhattan plots representing the statistical associations (on the y axis) between individual SNPs across the genome (on the x axis) and serum survival. The dotted red line represents the uncorrected significance threshold, and the dotted blue line represents the Sidak corrected (for multiple comparisons) threshold.

As the genome sequence for each of the 300 clinical *S. aureus* isolates was available we performed a GWAS (genome wide association study) to identify polymorphic loci that associated with the level of susceptibility of isolates to serum. For this, the data from the two distinct clones were analysed independently, with population structure within the clones being accounted for (Fig. 1c & d, Tables 1 & 2). We applied both uncorrected and corrected (for multiple comparisons) significance thresholds to this analysis, as our previous work has demonstrated that the stringency of multiple correction approaches increases the likelihood of type II errors, or false negative results. For the CC22 collection 12 loci were associated with serum survival and for the CC30s there were 31 loci associated (Fig. 1c & d, Tables 1 & 2).

**Tables 1 & 2:** Loci associated with serum survival in the CC22 and CC30 collections. The SNP position is relative to the origin of replication in the reference genomes: HO 5096 0412 (CC22 and MRSA252 (CC30). Locus tags and where available gene names or putative protein functions have been provided. NTML refers to the mutant available in the Nebraska transposon library.

| SNP position | CC22 locus tag | Description | -Log10(P) | NTML |
| --- | --- | --- | --- | --- |
| 1945336 | SAEMRSA15_RS09605 | Intergenic | 6.851546 | N/A |
| 375179 | SAEMRSA15_03110 | *metE* | 5.606418 | NE944 |
| 728154 | SAEMRSA15_06310 | *saeS* | 5.553331 | NE1296 |
| 1053036 | N/A | Intergenic | 3.60927 | N/A |
| 219764 | SAEMRSA15_01760 | putative oxidoreductase | 3.575929 | NE312 |
| 2280500 | SAEMRSA15_20910 | alcohol dehydrogenase | 2.257841 | NE1147 |
| 1196127 | SAEMRSA15_10750 | *yfhO* | 2.212486 | NE1597 |
| 408213 | SAEMRSA15_03430 | *guaA* | 1.876101 | N/A |
| 2492381 | SAEMRSA15_23120 | *tcyB* | 1.645759 | NE228 |
| 1514441 | SAEMRSA15_13420 | hypothetical protein | 1.600037 | NE1349 |
| 2430758 | SAEMRSA15_22540 | *tcaA* | 1.572702 | NE285 |
| 134276 | SAEMRSA15_01100 | hypothetical protein | 1.358386 | NE599 |

| SNP position | CC30 locus tag | Description | -Log10(P) | NTML |
| --- | --- | --- | --- | --- |
| 101174 | SAR0097 | putative DNA binding protein | 18.466 | N/A |
| 1188605 | SAR1143 | *arcC* | 12.786 | NE1719 |
| 900409 | SAR0856 | phosphoglycerate mutase | 11.226 | NE1422 |
| 2667418 | SAR2584 | *gntR* | 10.263 | NE1124 |
| 2421610 | SAR2347 | putative membrane protein | 10.160 | N/A |
| 1687821 | SAR1615 | *aroK* | 9.793 | N/A |
| 532944 | SAR0495 | *veg* | 9.751 | NE1703 |
| 1252982 | SAR1203 | *recG* | 8.502 | NE1344 |
| 1816815 | SAR1754 | *clpX* | 8.064 | N/A |
| 2206190 | SAR2143 | *ilvC* | 8.050 | NE1177 |
| 1443567 | SAR1386 | *trpA* | 6.647 | NE304 |
| 1167449 | SAR1119 | *uvrC* | 6.439 | NE1212 |
| 2442306 | SAR2374 | *ureC* | 6.425 | NE410 |
| 1784580 | SAR1719 | *tgt* | 6.396 | NE1885 |
| 880082 | SAR0835 | *est* | 6.354 | NE1122 |
| 1116463 | SAR1070 | *pdhD* | 5.600 | NE1610 |
| 289265 | SAR0248 | *tagB* | 5.324 | NE362 |
| 1442353 | SAR1385 | *trpB* | 4.910 | NE310 |
| 364861 | SAR0319 | NADH flavin oxidoreductase | 4.373 | N/A |
| 1963781 | SAR1877 | *menE* | 4.315 | N/A |
| 1946567 | SAR1856 | *arsB2* | 4.122 | NE1484 |
| 1416353 | SAR1364 | iron-sulfur cluster biosynthesis protein | 2.829 | NE1910 |
| 1887414 | SAR1810 | *fhs* | 2.741 | NE706 |
| 1133886 | SAR0251 | *tarF* | 2.725 | N/A |
| 30969 | SAR0023 | *sasH* | 2.507 | NE295 |
| 330358 | SAR0284 | *essC* | 2.486 | NE1484 |
| 2220348 | SAR2151 | *tex* | 1.704 | NE1079 |
| 1468392 | SAR1410 | *hipO* | 1.569 | NE1688 |
| 1652613 | SAR1582 | *zwf* | 1.556 | N/A |
| 2744774 | SAR2656 | *nrmA* | 1.549 | NE1223 |
| 2370812 | SAR2287 | *lacR* | 1.392 | NE346 |

Of the 43 loci associated with serum survival, there were transposon mutants available for 32 in the Nebraska library. To functionally verify our GWAS findings, each of these mutants were tested for their ability to survive exposure to human serum where we found seven mutants were significantly affected, and this effect was complemented by expressing the gene *in trans* (Fig. 2). The seven genes were *tcaA,* a gene associated with resistance to the antibiotic teicoplanin^23^; *tarK,* a gene involved in wall teichoic acid (WTA) biosynthesis^24^; *gntR,* which encodes gluconate kinase^25^; *ilvC,* which encodes ketol-acid reductoisomerase^26^; *arsB,* which is an efflux pump involved in arsenic resistance^27^; *yfhO,* which is involved in the glycosylation of lipoteichoic acid^28^; and *pdhD,* which encodes a lipoamide dehydrogenase^29^.

**Figure 2:**
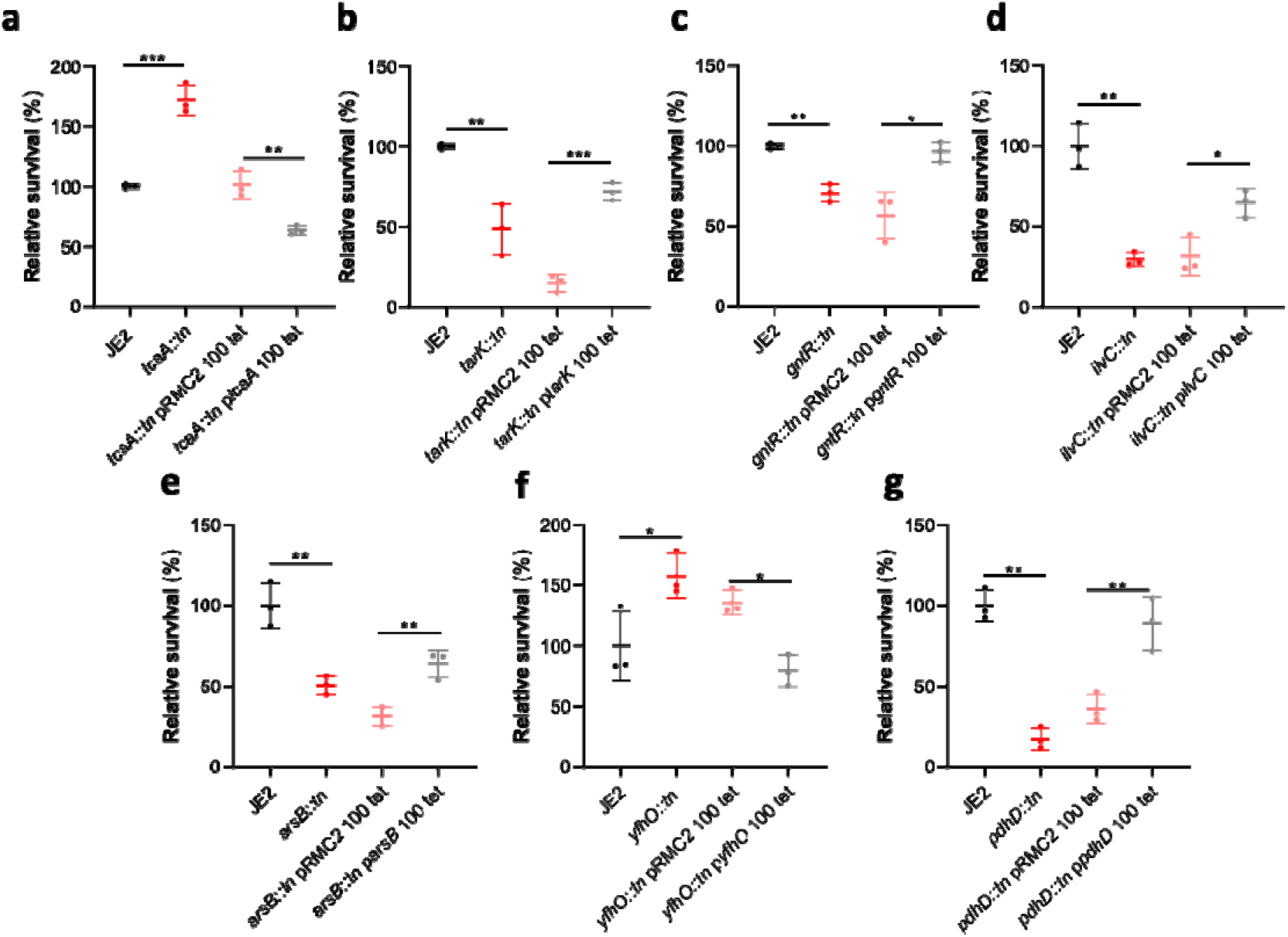
Functional verification of loci involved in serum sensitivity of *S. aureus.* The effect of the inactivation of the genes associated serum sensitivity was examined using transposon mutants. Of the 32 mutants tested seven were significantly affected: (**a**) *tcaA,* (**b**) *tarK,* (**c**) *gntK,* (**d**) *ilvC,* (**e**) *arsB,* (**f**) *yhfO* and (**g**) *pdhD.* The effect of the mutations on serum sensitivity was complemented by expressing the gene from the expression plasmid pRMC2. The dots represent individual data points, the bars the mean value, and the error bars the standard deviation. Significance was determined as * <0.05, ** <0.01, *** <0.001

To determine which aspects of human serum the identified mutants were either more or less sensitive to, we measured their relative ability to survive exposure to some of the anti-bacterial factors found in serum: to the HDFA arachidonic acid, to hydrogen peroxide and to two AMPs: HNP-1 and LL37 (Fig. 3). The TcaA and IlvC mutants were less sensitive to arachidonic acid when compared to the wild type strain (JE2), but no difference in the sensitivity of the other mutants to this HDFA was observed (Fig. 3a). The PdhD mutant was more sensitive to hydrogen peroxide, but the other mutants were all as sensitive as the wild type strain (Fig. 3b). The TcaA mutant was less sensitive to both HNP-1 and LL37, whereas the TarK, GntK, ArsB and PhdP mutants were all more sensitive to these AMPs (Fig. 3c & 3d).

**Figure 3:**
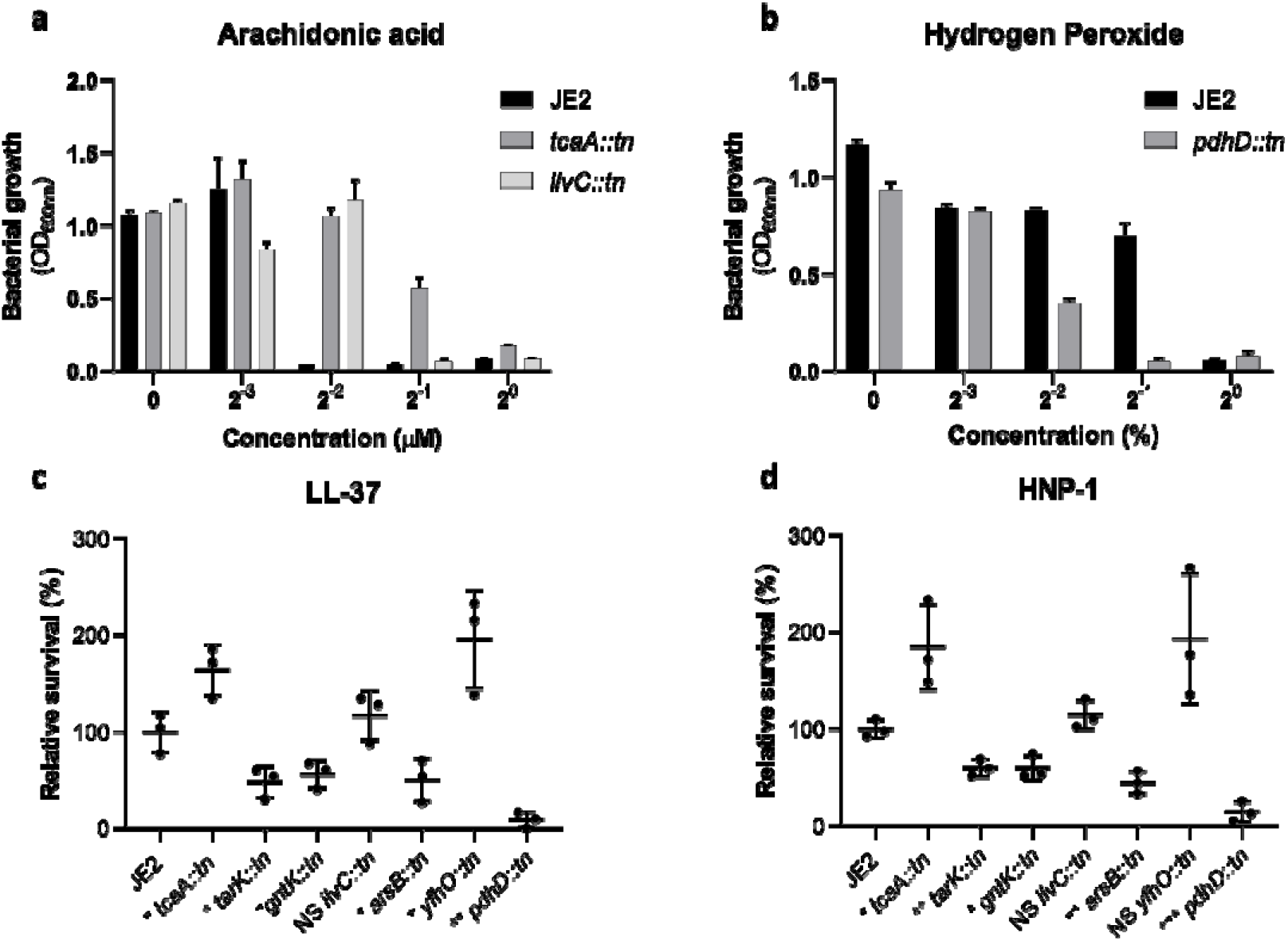
The serum sensitivity mutants display varying sensitivities to individual components of serum. (**a** & **b**) Growth of each mutant in 2-fold dilutions of a starting concentration of 400μM arachidonic acid and 0.1% hydrogen peroxide was quantified. The data provided represents growth measured by optical density (OD_600nm_) after 24h incubation. (**c** & **d**) The ability of the mutants to survive exposure to 5 μg/ml of LL-37 and 5 μg/ml of HNP-1 was quantified. The dots represent individual data points, the bars the mean value, and the error bars the standard deviation. Significance was determined as * <0.05, ** <0.01, *** <0.001.

The TcaA mutant is of particular interest as it was resistant to three of the four antimicrobial components of serum tested (Fig. 3). This suggests that during bacteraemia a TcaA mutant may cause a more severe bacteraemia relative to the wild type strain as it will survive in blood for longer. However, during bacteraemia the bacterial cells are exposed to whole blood as opposed to serum alone. To examine whether any of the other components of blood interferes with the decrease in sensitivity we have observed for the TcaA mutant, we repeated the sensitivity assay in both serum and whole blood, and we found the TcaA mutant to be less sensitivity to both, relative to the wild type strain (Fig. 4a). Murine models of *S. aureus* bacteraemia have utility when examining the activity of bacterial features that interact with host components common to both humans and the animal model, for example FnBPA, MspA and PurR^30–32^. However, significant differences in the types of HDFAs and AMPS contained in the serum of humans and mice exist. To examine whether a murine model of bacteraemia might be suitable to test whether a TcaA mutant displays increased virulence, we tested its relative sensitivity to mouse serum and found no difference relative to the wild type strain (Fig. 4B). Consistent with this in a murine model of *S. aureus* bacteraemia the TcaA mutant did not display any increase in virulence over the wild-type strains (Supp. Fig. 1). This suggests that the TcaA mediated sensitivity of *S. aureus* to serum is human specific.

**Fig. 4:**
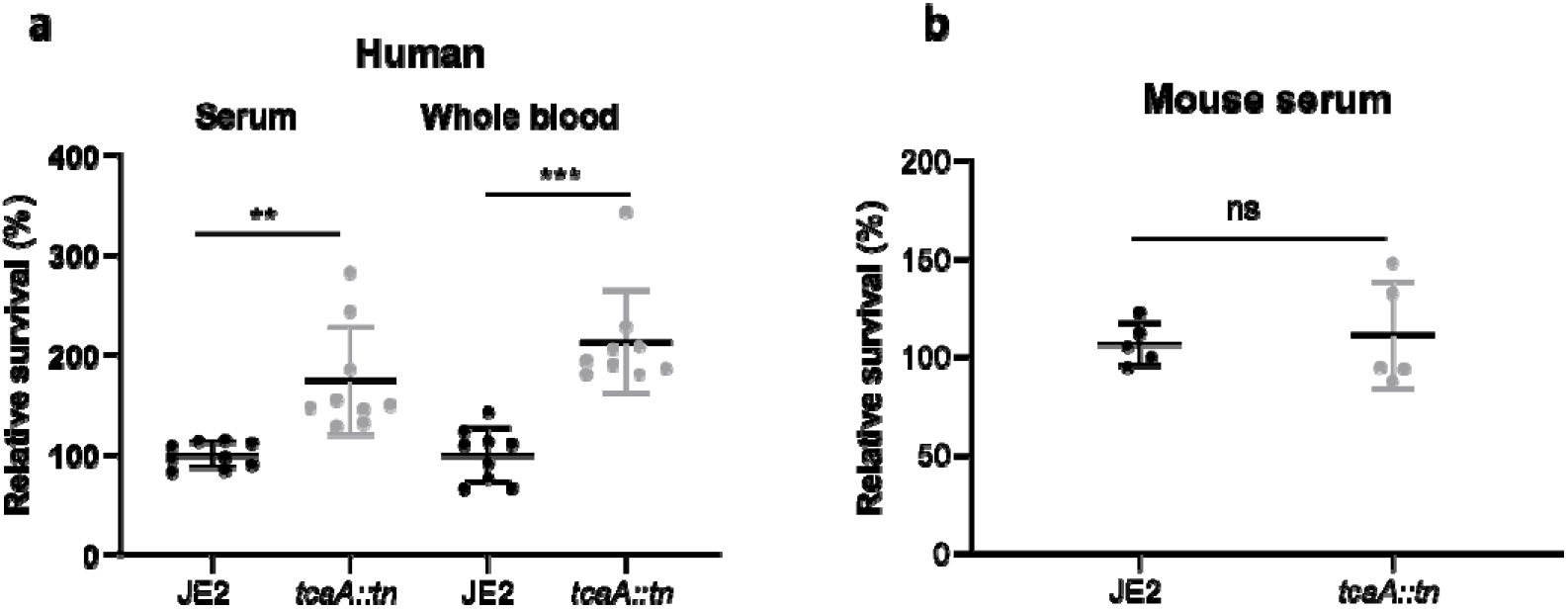
The decreased sensitivity of the TcaA mutant is specific to human serum. (**a**) The wild type *S. aureus* strain JE2 and the isogenic TcaA mutant were incubated in either serum of whole blood of human origin, and in both environments, the TcaA mutant survived better. (**b**) The wild type and TcaA mutants were exposed to mouse serum where no difference in survival between the bacteria was observed. The dots represent individual data points, the bars the mean value, and the error bars the standard deviation. Significance was determined as * <0.05, ** <0.01, *** <0.001.

Within our clinical strain collection there were 10 isolates with a single nucleotide polymorphism (SNP) in the *tcaA* gene, which conferred a Phe (290) to Ser change in the protein sequence. We have mapped the position of these isolates onto a figure displaying the range of serum sensitivities of the collection to visualise the signal detected by the GWAS. The majority of the isolates with the SNP were less sensitive to serum, suggesting this polymorphism negatively affected the activity of the protein (Fig. 5a). To understand the clinical significance of this, given the specificity of the activity of TcaA to human serum, we extended our analysis to a larger collection where the source of the isolate was recorded (i.e. bacteraemia, SSTI or carriage). We examined the distribution of nonsynonymous mutations in the *tcaA* gene across these, where it was present in 5.7% of the bacteraemia compared with only 1.4% of non-bacteraemia isolates (n= 20/348 and 2/139 respectively; p=0.068 chi-squared test). That isolates with defective *tcaA* genes are enriched amongst those causing bacteraemia supports our hypothesis that TcaA can limit *S. aureus’s* propensity to cause a bloodstream infection.

**Fig. 5:**
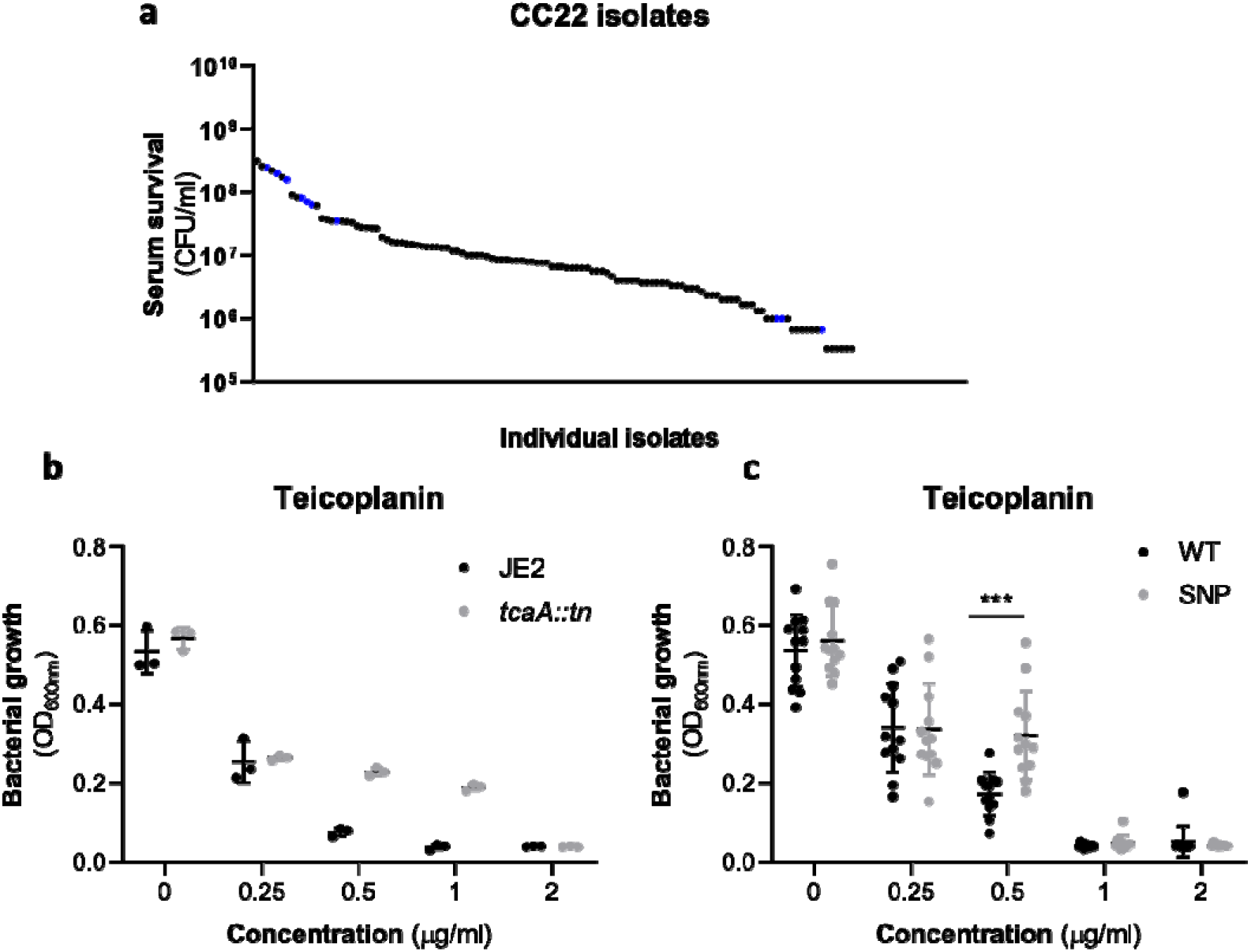
Polymorphism in the *tcaA* gene associate with both increased serum survival and increased teicoplanin resistance. (**a**) The individual clinical isolates with polymorphism in the *tcaA* gene are indicated (in blue) on a graph displaying the range of serum survival of the collection. (**b**) Growth of the wildtype and TcaA mutant in increasing concentrations of teicoplanin demonstrate that the mutant is less sensitive to this antibiotic. (**c**) The 10 clinical isolates with polymorphisms in the *tcaA* gene were on average more resistant than those with the wild type gene. The dots represent individual data points, the bars the mean value, and the error bars the standard deviation. Significance was determined as * <0.05, ** <0.01, *** <0.001.

An additional interesting feature to the TcaA protein is that is has also been associated with increased resistance to the antibiotic teicoplanin^23,33^. To verify that the inactivation of the *tcaA* gene in the JE2 background also affects teicoplanin resistance we quantified growth of the wild type and mutant in increasing concentrations of the antibiotic, where we found the effect to be consistent with previous literature (Fig. 5b). To examine whether the 10 clinical isolates with polymorphisms in *tcaA* were also less sensitive to teicoplanin, we compared their ability to grow in increasing concentrations of teicoplanin relative to 10 randomly selected isolates from the same collection with the wild type gene, i.e. without the polymorphism. The isolates with the polymorphism in the *tcaA* gene were on average less sensitive to teicoplanin (Fig. 5c).

The antibacterial properties of teicoplanin, HDFAs and AMPs are quite distinct. Teicoplanin is a glycopeptide antibiotic that inhibits the cross-linking of peptidoglycan layers^34^, whereas from what is known about the antibacterial mode of action of HDFAs and AMPs, they penetrate through the cell wall and attack the bacterial membrane^9–15^. AMPs are positively charged molecules that rely on the relatively negative charge across the bacterial cell envelope to penetrate, and resistance is frequently acquired by changing the charge across the cell wall such that the AMPs are repelled^14,15^. Less is known about how HDFAs penetrate the cell wall; however, resistance is associated with changes in the abundance of wall teichoic acids (WTA) in the bacterial cell walls^16^. WTA are hydrophilic and when present in the cell wall they are believed to be protective by interfering with the penetration of the hydrophobic HDFAs through to the bacterial membrane^16^. It has also been shown that when the ligation of WTA to the cell wall is affected, such that they are instead released into the environment, this also decreases the sensitivity of the bacteria to HDFAs^11^. With distinct targets and means of accessing these targets, it is intriguing to consider how TcaA may be contributing to increasing the sensitivity of *S. aureus* to all of three of these anti-bacterial molecules.

To examine whether the decreased sensitivity of the TcaA mutant to arachidonic acid is a result of an increase in the abundance of WTA in the bacterial cell we extracted and quantified WTA from the wild type and TcaA mutant, where we found contrary to our initial expectations that the cell wall in the TcaA mutant had significantly less WTA when compared to the wild type cell extract (Fig. 6a). With less WTA in the cell wall and recent work suggesting that when the WTA is released from the bacteria it decreases the sensitivity of *S. aureus* to arachidonic acid^11^, we quantified WTA in the bacterial supernatant of the wild type and TcaA mutant. There we found significantly more WTA in the TcaA mutant extract (Fig. 6b), which suggests that the TcaA mutant does not ligate WTA into the cell wall efficiently, and instead releases it from the cell wall. It is unclear from previous work how released WTA affects arachidonic acid sensitivity, and our hypothesis is that it can sequester it from the environment, thereby protecting the bacteria. To test this, we grew and harvested the supernatant of both the wild type and TcaA mutant, where the supernatant of the mutant will contain higher levels of released WTA. A culture of the wild type strain was pelleted, resuspended in one of the two supernatants and its ability to grow in increasing concentrations of arachidonic acid quantified. The supernatant of the TcaA mutant protected the wild type strain from the antibacterial action of arachidonic acid, which supports our hypothesis that the released WTA sequesters or inactivates the arachidonic acid (Fig. 6c), and explains why the mutant is less sensitive to this HDFA.

**Figure 6:**
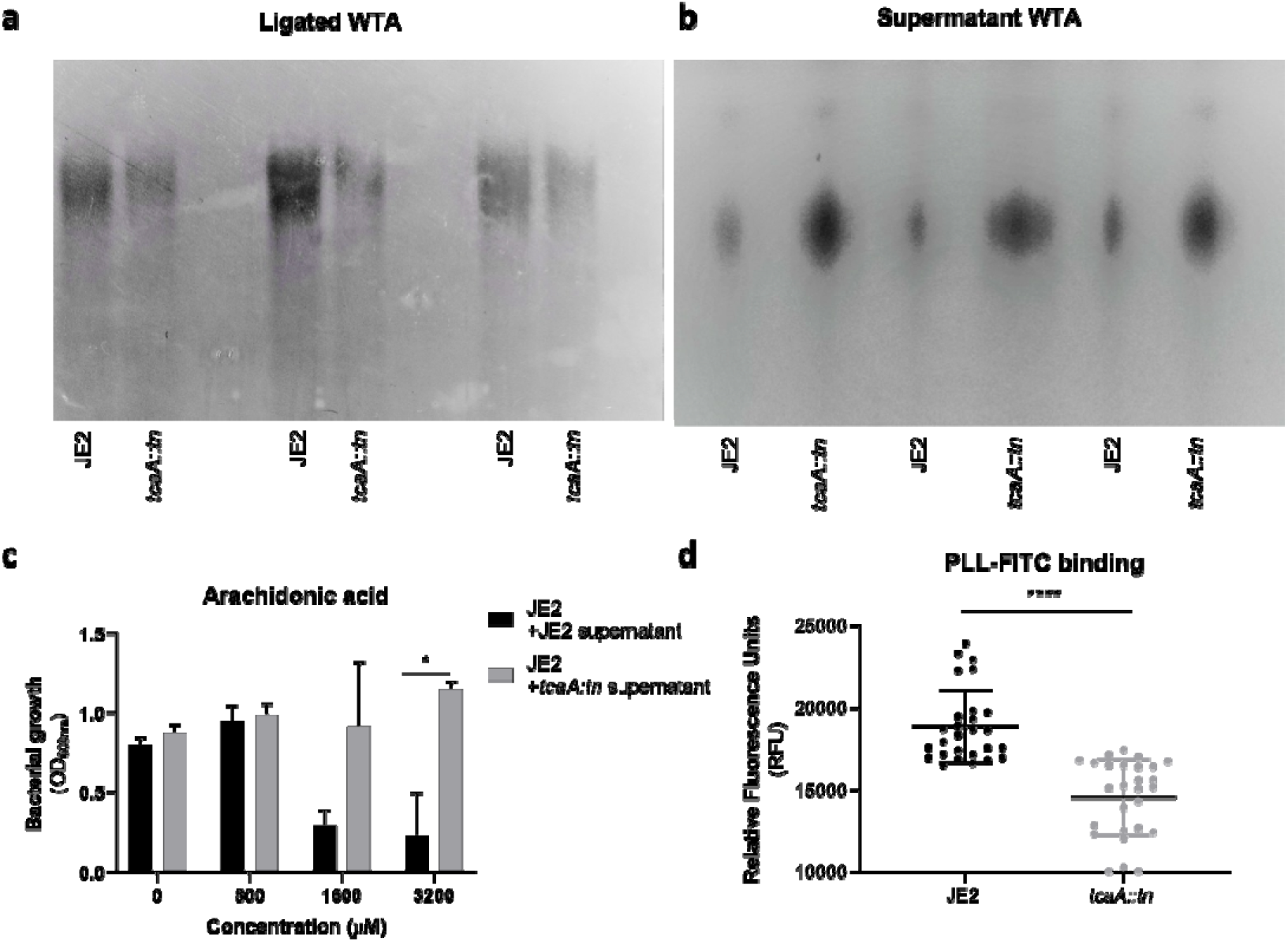
Wall teichoic acids are released from the cell wall in the TcaA mutant. (**a** & **b**) WTA was extracted from both cells and supernatant of the wild type and TcaA mutant of *S. aureus* and visualised on and SDS-PGAE gel stained with 1 mg/ml Alcian blue. The TcaA mutant had significantly less WTA in the cell wall but more in the supernatant. (**c**) The supernatant of JE2 and the TcaA mutant were harvested and a culture of JE2 resuspended in each. The relative ability of the JE2 to grow in these supernatants with increasing concentrations of arachidonic acid suggests that the released WTA in the mutant supernatant sequesters the arachidonic acid, thereby protecting the bacteria from being killed by the HDFA. (**d**) The charge across the cell wall of the wild type and TcaA mutant was compared using fluorescently labelled poly-L-lysine (PLL-FITC), where the mutant was found to be less negatively charged. The dots represent individual data points, the bars the mean value, and the error bars the standard deviation. Significance was determined as * <0.05, ** <0.01, *** <0.001, **** <0.0001.

In additional to being hydrophilic, WTA is also negatively charged^24^. Given that a change in charge across the bacterial cell wall is frequently associated with resistance to AMPs^13,15^, and that there is less WTA in the cell wall of the TcaA mutant, we hypothesised that a change in charge as a result of the reduced WTA may explain the decreased AMP sensitivity. To examine this, we incubated the bacteria with fluorescently labelled poly-L-lysine (PLL-FITC), which is positively charged and through its charge-related ability binds to and fluorescently label cells, providing a measure of the charge across bacterial cell walls. Using this we found that the TcaA mutant bound significantly less PLL-FITC and is therefore less negatively charged than the wild type strain, which explains the decrease in AMP sensitivity we have found associated with this mutant (Fig. 6d). These results explain why the TcaA mutant is less sensitive to both fatty acids and AMPs. The release of WTA sequesters the fatty acids from the environment, and due to their negative charge, the loss of WTA in the cell wall affects the charge across the cell wall and results in the repulsion of the positively charged AMPs. What role the change in ligation of WTA to the cell wall plays in resistance to teicoplanin is as still unclear and is currently under investigation.

As TcaA has been shown to be upregulated when exposed to teicoplanin^23,33^, we examined whether serum would also induce its expression. We exposed the wild type bacteria to either serum (10%) or teicoplanin (10μg/ml) for 5, 20 and 90 minutes, our expectation being that at 90 minutes most of the bacterial cells will have died. We extracted the mRNA and performed qRT-PCR to quantify the level of transcription of the *tcaA* gene as well as the *gyrB* housekeeping gene. Both teicoplanin and serum induced a rapid increase in *tcaA* transcription (Fig. 7a). To examine whether this increased expression would result in cross resistance we again exposed the wild type bacteria to either serum or teicoplanin, but this time at subinhibitory concentrations. The teicoplanin exposed bacteria were then incubated in a subinhibitory concentration of serum (fig. 7b), and the serum exposed bacteria incubated in subinhibitory concentrations of teicoplanin (fig. 7c). Their relative ability to survive this secondary exposure having had *tcaA* expression induced by the first exposure was quantified. In both cases pre-exposure to either serum or teicoplanin was sufficient to induce increased sensitivity to the other (Fig. 7b and 5c). These results suggest that the induction of expression of *tcaA* upon exposure to either teicoplanin or serum leads to an increase in sensitivity to both of these antibacterial agents.

**Figure 7:**
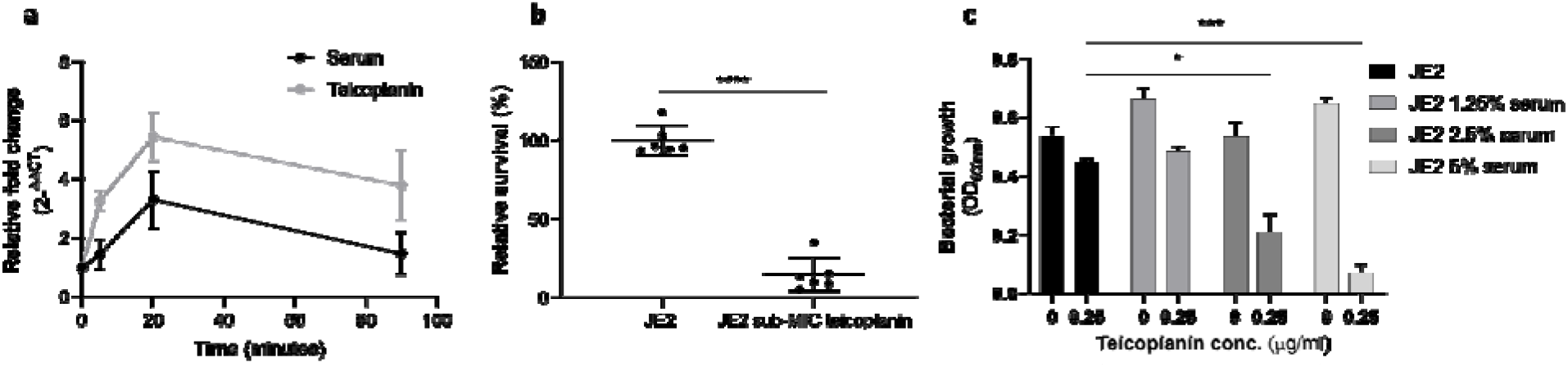
TcaA expression is induced upon exposure to both teicoplanin and serum. (**a**) The expression of the *tcaA* gene was quantified by qRT-PCR following 5, 20 and 90 mins exposure to both human serum (10%) and teicoplanin (10 μg/ml), and both exposures induced an significant increase in transcription. (**b**) Serum survival assays were performed where the wild type *S. aureus* strain JE2 was pre-exposed to sub-inhibitory concentrations (0.25 μg/ml) of teicoplanin, which significantly increased its sensitivity to serum killing. (**c**) Growth of JE2 in the presence of sub-inhibitory concentrations of teicoplanin (0.25 μg/ml) over a 24h period was quantified following pre-exposure to increasing concentrations of human serum (1.25, 2.5 and 5%), where preexposure to serum increased sensitivity to teicoplanin. The dots represent individual data points, the bars the mean value, and the error bars the standard deviation. Significance was determined as * <0.05, ** <0.01, *** <0.001, **** <0.0001.

## Discussion

To develop novel therapeutic approaches for infectious diseases, we need to understand how microorganisms cause disease and evade the host’s immune system. To address this, we have applied a population-based approach to analyse the pathogenicity of *S. aureus* and identified seven genes that affect the bacteria’s ability to survive exposure to human serum, the first step in the development of bacteraemia. The dissection of the molecular detail of how these genes affect serum survival is underway to determine their potential for therapeutic intervention. However, our focus here has been on the *tcaA* gene where we show that its expression is induced upon exposure to serum, that it is involved in the ligation of WTA into the cell wall, and that this renders the bacteria more susceptible to both AMPs and HDFAs (a graphical summary has been provided Fig. 8). It is interesting to note that *tarK,* which is one of the other six genes associated with serum survival is involved in the biosynthesis of WTA, providing further evidence for the importance of these cell wall associated molecules to this aspect of the biology and pathogenicity of *S. aureus.*

**Figure 8:**
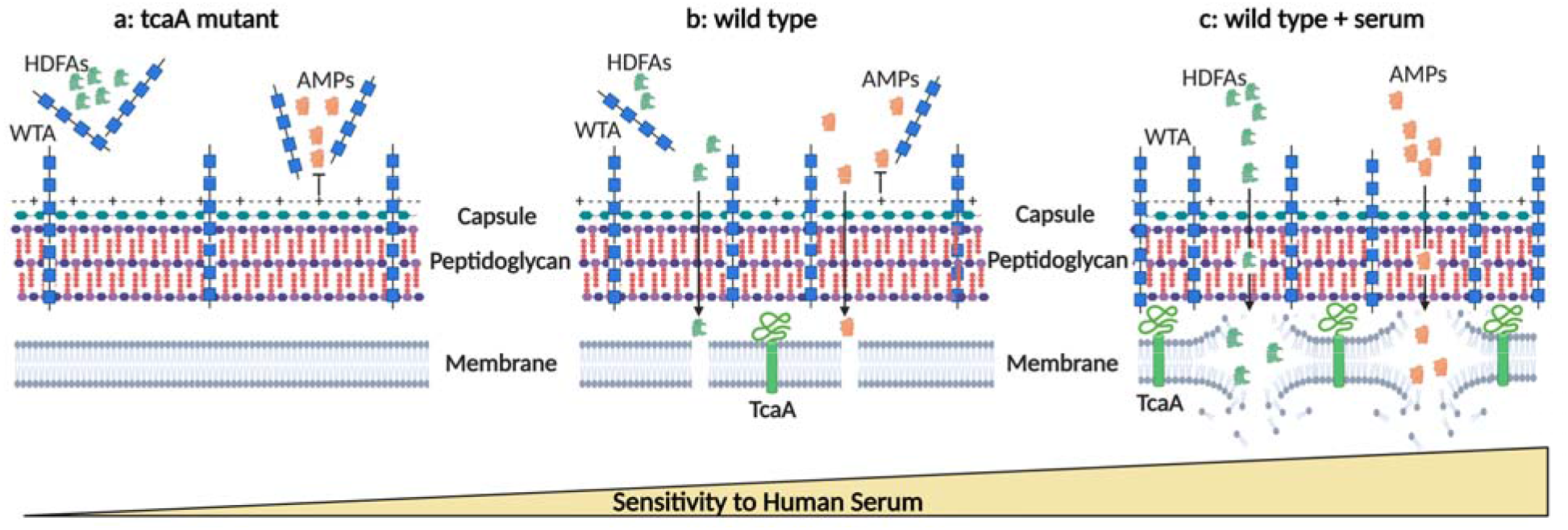
Graphical summary of TcaA activity. (**a**) The abundance of WTA in the TcaA mutant is relatively low within the cell wall, and relatively high in the extracellular milieu. This increases the positive charge across the bacterial surface which repels the positively charged AMPs, increasing the bacteria’s resistance to these. With higher levels of WTA in the environment this facilitated greater levels of sequestration of the AFAs resulting in increased resistance of the mutant bacteria to HDFAs. (**b**) The wild type *S. aureus* produces a moderate level of TcaA which results in moderate levels of WTA in both the cell wall and the extracellular milieu. Due to WTA’s negative charge this results in charge across the bacterial surface remaining relatively negative, which allows some of the AMPs to penetrate through and attack the cell wall. The release of some WTAs into the external environment will sequester some HDFAs, increasing the resistance of the bacteria to these. (**c**) Upon exposure to serum the expression of TcaA in the wild type strain is induced which will result in high levels of WTA in the cell wall and low levels in the extracellular milieu. The relative decrease in the negativity of the charge across the bacterial surface this causes renders the bacteria more susceptible to AMPs. With WTA securely ligated to the cell wall there are none available in the extracellular milieu to sequester the HDFAs, rendering the bacteria more susceptible to membrane disruption by these molecules. Image created using BioRender.

The evolution of the virulence of a pathogen is heavily dependent upon their mode of transmission, which for opportunistic pathogens raises interesting challenges. Increased invasiveness facilitates enhanced within-host fitness, as the bacteria multiply and spread throughout the body. However, if invasiveness limits between-host fitness by preventing them from transmitting to a new host, how do opportunistic members of our microbiome limit or control their invasiveness to enhance their long-term success? For *S. aureus,* as a classic example of an opportunist, entry into the bloodstream has deleterious consequences for both the host and pathogen, which begs the question as to why bacterial features that facilitate this have been selected for. To understand this we need to consider the other ways in which *S. aureus* interact with humans: as both a commensal and as a major cause of SSTIs, from which it can readily transmit. So perhaps the protective role these genes play are also important during those interactions, which would confer a selective advantage, and it is just an unfortunate coincidence that they also confers increased survival in serum during the development of bacteraemia. In support of the argument that the ability of *S. aureus* to cause bacteraemia should be selected against, we identified two genes that contribute negatively to this, with one, TcaA, induced upon exposure to serum. What role this protein may play during other host interactions is yet to be determined, possibly involving the need for WTA for colonisation, but during bacteraemia it appears to play the role of a ‘suicide’ factor, limiting the ability of the bacteria to establish a bloodstream infection.

## Materials and Methods

### Bacterial Strains and Growth Conditions

A list of bacterial strains can be found in Table 3. All strains were cultured in Tryptic soy broth (TSB) for 18 h at 37°C with shaking. Nebraska transposon mutant library (NTML)^35^ mutants were selected for using erythromycin (5 μg/ml). For the complemented pRMC2^36^ strains, chloramphenicol (10 μg/ml) and anhydrous tetracycline (200 ng/ul) were added to the media where indicated.

**Table 3.** Strains used in this study.

| Strain | Description | Reference |
| --- | --- | --- |
| JE2 | USA300; CA -MRSA, type IV SCCmec; lacking plasmids p01 and p03; wild -type strain of the NTML | 35 |
| *tcaA:tn* | *tcaA* transposon mutant in JE2 | 35 |
| *tcaA:tn* pRMC2 | *tcaA* transposon mutant in JE2 transformed with empty pRMC2 vector | This study |
| *tcaA:tn ptcaA* | *tcaA* transposon mutant complemented with *tcaA* gene housed in pRMC2 expression plasmid | This study |
| *tarK:tn* | *tarK* transposon mutant in JE2 | 35 |
| *tarK:tn* pRMC2 | *tarK* transposon mutant in JE2 transformed with empty pRMC2 vector | This study |
| *tarK:tn ptarK* | *tarK* transposon mutant complemented with *tarK* gene housed in pRMC2 expression plasmid | This study |
| *gntR:tn* | *gntK* transposon mutant in JE2 | 35 |
| *gntR:tn* pRMC2 | *gntK* transposon mutant in JE2 transformed with empty pRMC2 vector | This study |
| *gntR:tn pgntR* | *gntK* transposon mutant complemented with *gntK* gene housed in pRMC2 expression plasmid | This study |
| *ilvC:tn* | *ilvC* transposon mutant in JE2 | 35 |
| *ilvC:tn* pRMC2 | *ilvC* transposon mutant in JE2 transformed with empty pRMC2 vector | This study |
| *ilvC:tn pilvC* | *ilvC* transposon mutant complemented with *ilvC* gene housed in pRMC2 expression plasmid | This study |
| *arsB:tn* | *arsB* transposon mutant in JE2 | 35 |
| *arsB:tn* pRMC2 | *arsB* transposon mutant in JE2 transformed with empty pRMC2 vector | This study |
| *arsB:tn parsB* | *arsB* transposon mutant complemented with *arsB* gene housed in pRMC2 expression plasmid | This study |
| *yfhO:tn* | *yfhO* transposon mutant in JE2 | 35 |
| *yfhO:tn* pRMC2 | *yfhO* transposon mutant in JE2 transformed with empty pRMC2 vector | This study |
| *yfhO:tn pyfhO* | *yfhO* transposon mutant complemented with *yfhO* gene housed in pRMC2 expression plasmid | This study |
| *pdhD:tn* | *pdhD* transposon mutant in JE2 | 35 |
| *pdhD:tn* pRMC2 | *pdhD* transposon mutant in JE2 transformed with empty pRMC2 vector | This study |
| *pdhD:tn ppdhD* | *pdhD* transposon mutant complemented with *pdhD* gene housed in pRMC2 expression plasmid | This study |

### Serum Resistance Assay

Overnight cultures with 0.25 μg/ml teicoplanin added where indicated were normalised to an OD_600nm_ of 0.1 and incubated in 10% pooled human serum (diluted in PBS) for 90 min at 37°C with shaking. Serial dilutions were plated on tryptic soy agar (TSA) to determine CFUs. The same number of bacterial cells inoculated into PBS, diluted, and plated acted as a control. Survival was determined as the percentage of CFU in serum relative to the PBS control. Relative survival was determined through normalisation to JE2.

### GWAS

Genome-wide association mapping was conducted using a generalized linear model, with capsule production as the quantitative response variable. We accounted for bacterial population substructure by adding to the regression model the first two component from a *principal component decomposition* of SNP data for each set of clinical samples (CC22 and CC30). The first two components accounted for 32% and 40% of the total variance for CC22 and CC30, respectively. In both cases, three distinct clusters were identified. We further considered a third model where we used cluster membership as covariates in our regression model, where clusters were defined using K-means clustering analysis (setting K=3); this, however, yielded identical results to the one based on PCA components. In total, 2066 (CC22) and 3189 (CC30) unique SNPs were analysed, the majority of which were subsequently filtered out for exhibiting a minor allele frequency (maf) of <0.03, reducing the data to 378 and 1124 SNPs, respectively. Reported P-values are not corrected for multiple comparison; Sidak corrected significance thresholds are indicated in the Manhattan plots.

### Genetic manipulation

Wildtype genes were amplified by PCR from JE2 genomic DNA using the primers shown in table 4 and KAPA HiFi polymerase (Roche). The PCR product was cloned into the tetracycline inducible plasmid pRMC2 using *Kpn*I and *Sac*I restriction sites and T4 DNA ligase (NEB). This was transformed into RN4220 and eventually into the respective NTML mutants through electroporation.

**Table 4.**
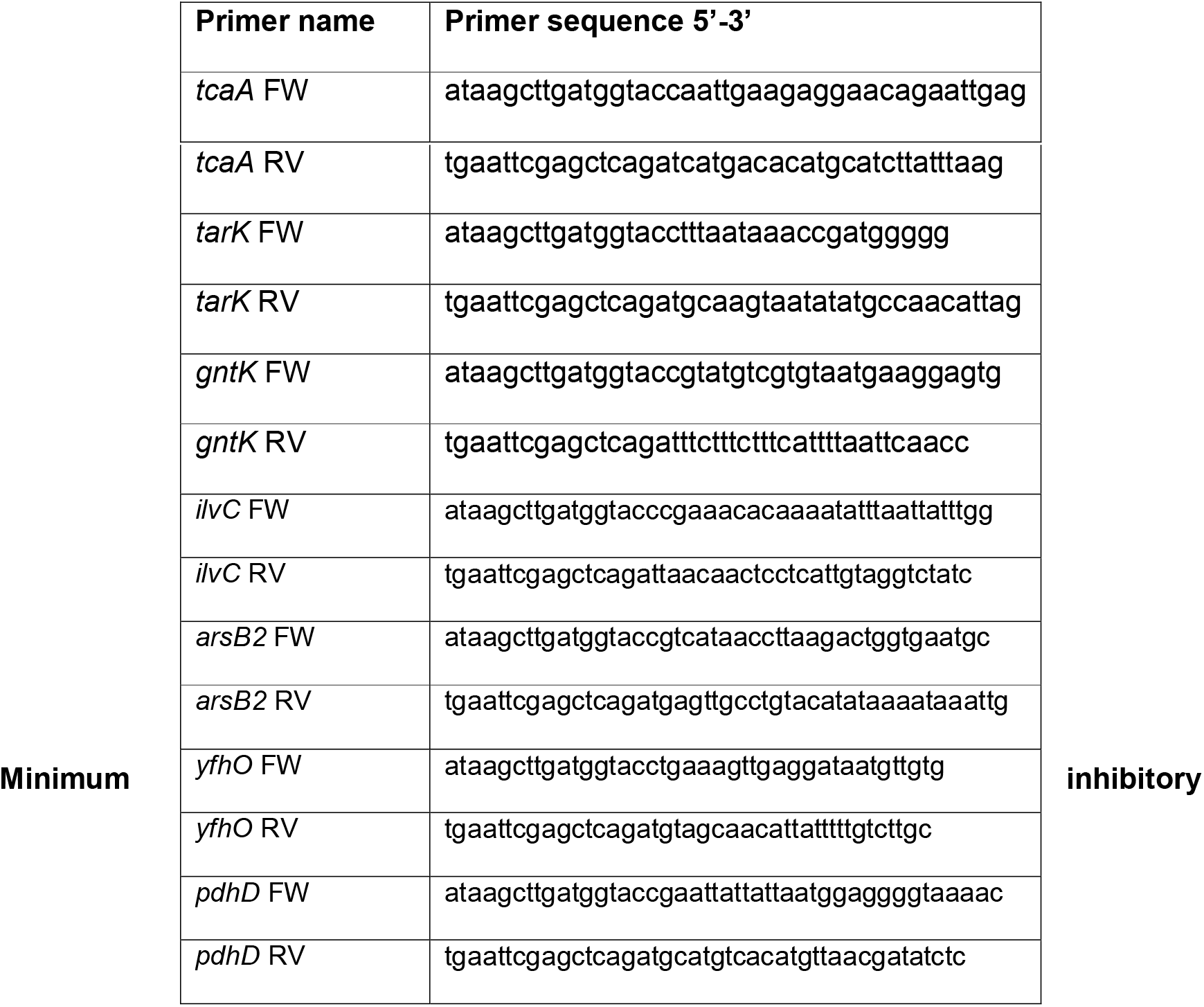
PCR primers used in this study.

### concentrations (MIC)

MICs were performed according to the micro broth dilution method^37^. Briefly, overnight cultures, with either 1.25%, 2.5% or 5% human serum added where indicated, were normalised to an OD_600nm_ of 0.1 in cation adjusted Mueller Hinton broth (MHB++) and 20 μl of resultant suspension used to inoculate 180 ul of fresh MHB++ containing either arachidonic acid (Sigma) or hydrogen peroxide (Sigma). 1:2 dilutions were subsequently performed and incubated for 20 h at 37°C without shaking. The ability of the bacteria to survive the antibiotics was determined by quantifying bacterial growth (OD_600nm_) using a CLARIOstar plate reader (BMG Labtech).

### Antimicrobial peptide susceptibility

Antimicrobial peptide susceptibility Human neutrophil defensin-1 (hNP-1) (AnaSpec Incorporated, California, USA) and LL-37 (Sigma) susceptibility assays were performed as described previously (1). Briefly, overnight cultures were normalised to an OD_600nm_ of 0.1 and incubated with 5 μg/mL of hNP-1 or LL-37 for 2 h at 37 °C. Serial dilutions were plated on tryptic soy agar (TSA) to determine CFUs. The same number of bacterial cells inoculated into PBS, diluted, and plated acted as a control. Survival was determined as the percentage of CFU in serum relative to the PBS control. Relative survival was determined through normalisation to JE2.

### WTA preparations

Crude WTA from murine sacculi was extracted for analysis by PAGE using adaptations of a previously described methodology^11^. Overnight cultures were washed once in buffer 1 (50 mM MES, pH 6.5) followed by centrifugation at 5,000g. Cells were resuspended in buffer 2 (4% [wt/vol] SDS, 50 mM MES, pH 6.5) and boiled for 1 hour. Sacculi were centrifuged at 5,000g and washed once in buffer 1, once in buffer 2, once in buffer 3 (2% NaCl, 50 mM MES, pH 6.5), once more in buffer 1 and finally resuspended in digestion buffer (20 mM Tris-HCl pH 8.0, 0.5% [w/v] SDS). To the digestion buffer suspension, 10 μl of proteinase K solution (2 mg/ml) was added and incubated on a heat block 50°C for 4 h at 1,400 rpm. Sacculi were centrifuged at 16,000 g and washed once in buffer 3, followed by 3 washes in dH_2_O to removed SDS. Sacculi were responded in 0.1M NaOH and incubated for 16 h at room temperature at 1,400 rpm. Following the incubation, the sacculi were centrifuged at 16,000g, leaving the teichoic acids in the supernatant. 250 μl of 1 M Tris-HCL pH 6.8 was added to neutralise the NaOH and stored at −20°C.

WTA was also precipitated from the supernatant of overnight culture by adding 3 volumes of 95% ethanol and incubation at 4°C for 2 hours^11^. Precipitated material was separated by centrifugation at 16,000 g for 15 min, washed once in 70% ethanol and resuspended in 100 mM Tris-HCL (pH 7,5) containing 5 mM CaCl_2_, 25 mM MgCl_2_, DNase (10 μg/ml) and RNase (50 μg/ml) and incubated for 3 h at 37°C. The enzymes were heat inactivated at 95°C for 3 min. The supernatant WTA preparations were similarly stored at −20°C before being analysed by PAGE.

### WTA PAGE

WTA preparations were separated on tricine polyacrylamide gels using a BioRad tetra cell according to a previously described method^11^. The gels were separated at 4°C using a constant amperage of 40 mA under constant stirring until the dye front reached the bottom. Gels were washed 3 times in MilliQ H_2_O followed by staining in 1 mg/ml Alcian blue overnight. Gels were subsequently destained in in MilliQ H_2_O, until the WTA became visible and finally imaged.

### Arachidonic acid supernatant conditioning

Overnight cultures of JE2 and the TcaA mutant were pelleted, and the supernatant saved. MICs were subsequently performed by adding 20 μl of OD_600nm_ 0.1 bacterial suspension to 50% overnight supernatant diluted in fresh MHB++ containing doubling concentrations of arachidonic acid (Sigma). The ability of the bacteria to survive the arachidonic acid was determined by quantifying bacterial growth (OD_600nm_) following 24h at 37°C using a CLARIOstar plate reader (BMG Labtech).

### FITC-Poly-L-lysine binding assay

A FITC-Poly L lysine binding assay was used to determine the relative surface charge of the bacteria (6). To measure FITC-PLL binding, 1 ml aliquots of *S. aureus* cultures (at OD_600nm_ 0.5 in PBS) were incubated with 80 μg/ml FITC-PLL (MW 15,000 – 30,000; Sigma) for 10 min at room temperature in the dark. Cells were then washed with PBS by three rounds of centrifugation (16000 × g for 1 min). Samples were stored in the dark. Fluorescence was visualised by using a CLARIOstar plate reader (excitation 485 nm: emission 525 nm).

### mRNA extraction

Overnight cultures were back-diluted to an OD_600nm_ of 0.05 in 50 ml of fresh TSB and grown to an OD6_00nm_ of 2. 200 μl of either PBS, 100% serum or 100 μg/ml teicoplanin diluted in PBS (Sigma) was added to 1.8 ml of bacterial culture and incubated for either 5, 20 or 90 min. RNA was extracted by Quick-RNA Fungal/Bacterial Miniprep Kit (Zymo Research) according to the manufacturer’s instructions. RNA integrity was checked by running 5 μL aliquot of the RNA on a 1% agarose gel and observing the intensity of the ribosomal RNA (rRNA). RNA samples were treated by TURBO^™^ DNase (Invitrogen) to eliminate any genomic DNA contamination. To verify that the samples were free from any DNA contamination, RNA samples were subjected to RT-qPCR alongside with a no template control (NTC) and 2.5 ng of a known genomic DNA and threshold rate were compared.

### Quantitative reverse transcriptase (RT-qPCR)

To quantify the expression of the *tcaA* gene, RT-qPCR was performed using the housekeeping *gyrB* gene as a control. Complementary DNA (cDNA) was generated from mRNA using qScript^®^ cDNA Synthesis Kit (Quantabio). Following the manufacturers protocol the cDNA was used as a template for the qPCR reaction. Primers used are listed in table 3. The reverse-transcriptase PCR (RT-PCR) was performed as follows: 10 μl 2x KAPA SYBR Mix, 1 μl of 10 μM forward primer, 1 μl of 10 μM μl reverse primer, 5 μl cDNA and RNase-free water up to a total of 20 μl volume. The PCR condition cycles consisted of initial denaturation at 95°C for 2 min followed by 35 cycles of denaturation at 95°C for 30 s, annealing at 55°C for 30 s and extension at 72°C for 30 s. RT-PCR was carried out in technical triplicate for each sample and 3 biological repeats. The ratio of *tcaA* and *gyrB* transcript number was calculated using the using the 2^−(ΔΔCt)^ method.

### In vivo intra-venous challenge model

C57/Bl6 mice were injected with 5X10^7^ CFU of JE2 or *tcaA::tn* via the tail vein. Mice were culled at specific time points post challenge and blood collected by cardiac puncture. Blood was diluted in sterile PBS and plated on TSA to determine CFU/ml.

### Ethics statement

C57/Bl6 mice were bred in-house in Trinity College Dublin. All mice were housed under specific pathogen-free conditions at the Trinity College Dublin Comparative Medicines unit. All mice were used at 6–8 weeks. All animal experiments were conducted in accordance with the recommendations and guidelines of the health product regulatory authority (HPRA), the competent authority in Ireland and in accordance with protocols approved by Trinity College Dublin Animal Research Ethics Committee.

### Statistics

Paired two-tailed student t-test or One-way ANOVA (GraphPad Prism v9.0) were used to analyse the observed differences between experimental results. A *p*-value <0.05 was considered statistically significant.

## Supplementary Figure.

**Supp. Fig. 1:**
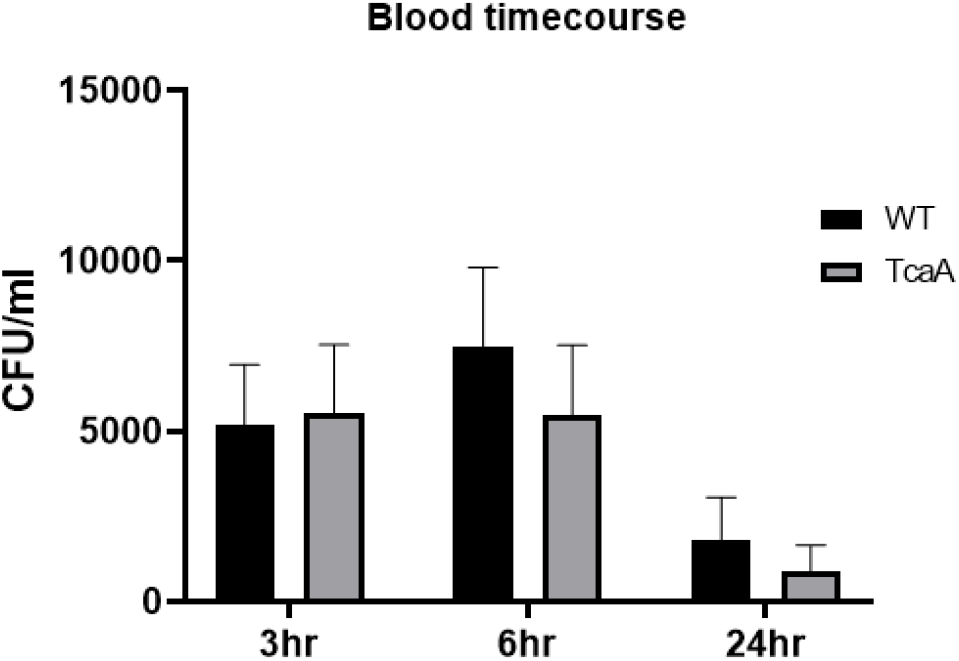
The TcaA mutant is not attenuated in a mouse model fo sepsis. C57/Bl6 mice were injected with 5X10^7^ CFU of JE2 or *tcaA::tn* via the tail vein. Mice were culled at specific time points post challenge and blood collected by cardiac puncture. Blood was diluted in sterile PBS and plated on TSA to determine CFU/ml.

